# Socially Integrated Polysubstance (SIP) system: an open-source solution for continuous monitoring of polysubstance fluid intake in group housed mice

**DOI:** 10.1101/2022.10.11.511835

**Authors:** Katrina Wong, Ziheng Christina Wang, Makenzie Patarino, Britahny Baskin, Suhjung Janet Lee, Abigail G. Schindler

## Abstract

Despite impressive results from neuroscience research using rodent models, there is a paucity of successful translation from preclinical findings to effective pharmacological interventions for treatment of substance use disorder (SUD) in humans. One potential reason for lack of translation from animal models is difficulty in accurately replicating the lived experience of people who use drugs. Aspects of substance use in humans that are often not modeled in animal research include but are not limited to 1) voluntary timing and frequency of substance intake, 2) social environment during substance use, and 3) access to multiple substances and multiple concentrations of each substance. Critically, existing commercial equipment that allows for social housing and voluntary polysubstance use (e.g., home cage monitoring system) is prohibitively expensive and no open-source solutions exist. With these goals in mind, here we detail development of the Socially Integrated Polysubstance (SIP) system, an open-source and lower cost solution that allows for group housed rodents to self-administer multiple substances with continuous monitoring and measurement. In our current setup, each SIP cage contains four drinking stations, and each station is equipped with a RFID sensor and sipper tube connected to a unique fluid reservoir. Using this system, we can track which animal (implanted with unique RFID transponder) visits which drinking location and the amount they drink during each visit (in 20 ul increments). Using four flavors of Kool-Aid, here we demonstrate that the SIP system is reliable and accurate with high temporal resolution for long term monitoring of substance intake and behavior tracking in a social environment. The SIP cage system is a first step towards designing an accessible and flexible rodent model of substance use that more closely resembles the experience of people who use drugs.

## Introduction

Substance use and addiction research has evolved significantly over the years, and it is now generally accepted that substance use disorders (SUDs) are disorders involving the brain that have complex biopsychosocial etiology (Strickland et al., 2022; Stull et al., 2022; Venniro et al., 2020). Despite impressive results from preclinical neuroscience research using rodent models, few new effective treatments have been successfully translated to the clinic. This has led to questioning whether animal models are a valid method to study SUDs, and increasing emphasis placed on improving ethological relevance and connection to lived experience of people who use drugs (Ahmed et al., 2013; Field & Kersbergen, 2020; Negus & Banks, 2020; Strickland et al., 2022; Stull et al., 2022; Vandaele & Daeppen, 2022; Venniro et al., 2020). Specifically, features of the human condition that can be, but are not often, addressed using rodent models are: 1) voluntary timing and frequency of substance intake, 2) social environment during substance use, and 3) access to multiple substances and multiple concentrations of the same substance (including food and water as alternative reinforcers). Critically, current commercially available options that provide these features (e.g., IntelliCage) are prohibitively expensive. As a lower-cost alternative, here we detail development of the Socially Integrated Polysubstance (SIP) system, an open-source solution that allows for group-housed rodents to self-administer multiple substances with continuous monitoring and measurement.

The human population displays a wide variety of individual differences in response to rewarding substances and a majority of people who use drugs do not go on to develop “problematic” use or diagnosis of SUD (Strickland et al., 2022; Stull et al., 2022; Vandaele & Daeppen, 2022; Venniro et al., 2020). The 2020 National Survey on Drug Use and Health reported SUD rates in respondents aged 12 and older between 1.0% (any opioid) and 10.2% (alcohol) (SAMHSA, 2020). While these percentages are relatively low, they still translate to over 17 million people with past month heavy alcohol use (5 or more binge drinking episodes in the past 30 days) and over 40 million people with any past year SUD diagnosis. Critically, this report and others also highlight high rates of polysubstance use (use of multiple substances) (Compton et al., 2021; Crummy et al., 2020; Hazani et al., 2022; James et al., 2021; SAMHSA, 2020). Polysubstance use can include use changes across the lifespan (longitudinal), sequential (use of one substance followed by use of another substance), and simultaneous co-use of substances (use of multiple substances at the same time). While polysubstance use has received increased attention in the animal literature recently, these reports almost exclusively rely on longitudinal (e.g., one substance during adolescence and a different substance during adulthood), sequential (separate use in different self-administration sessions), or forced simultaneous use (self-administration of a mix of substances in the same solution) (Amico et al., 2022; Doyle et al., 2022; James et al., 2021; O’Rourke et al., 2016; Pattison et al., 2012; Stennett et al., 2020), with a minority investigating simultaneous voluntary use of multiple substances (Amico et al., 2022; Foo et al., 2022; O’Rourke et al., 2016; Vandaele et al., 2022).

When trying to model the human condition of substance use and development of SUD, it is important to note that a variety of substances, in a variety of concentrations, are often available (e.g., polysubstance), in addition to a multitude of alternative reinforcers such as food and social interaction. Indeed, there has been increasing focus placed on using drug-choice procedures to study substance use and model aspects of SUD in rodents (Ahmed et al., 2013; Strickland et al., 2022; Venniro et al., 2020), with numerous studies now demonstrating that rodents often choose food or social reward over drugs such as methamphetamine or heroin (Cantin et al., 2010; Caprioli et al., 2015; Huynh et al., 2017; Kearns et al., 2017; Lenoir et al., 2007; Lenoir & Ahmed, 2008; Russo et al., 2018; Venniro et al., 2018). Likewise, a recent study demonstrated that volitional control over substance choice (between cocaine and saccharin) completely abolished cocaine choice in a majority of rats (Vandaele et al., 2022). Finally, in a four bottle choice procedure (3 does of ethanol and water), ethanol dose preference was predictive of subsequent drinking behavior when the ethanol solutions were devalued by addition of the bitter adulterant quinine (Foo et al., 2022). In addition to availability and type of reinforcer, animal housing environment (group housed vs. socially isolated; enriched vs. deprived homecage environment) can drastically impact outcomes related to substance intake and preference (Ahmed et al., 1995; Althobaiti & Almalki, 2020; Engeln et al., 2021; Famitafreshi et al., 2016; Holgate et al., 2017; Karkhanis et al., 2019; E. D. Klein et al., 2007; Mastrogiovanni et al., 2021; Nyuyki et al., 2012; Panksepp et al., 2017; Scott et al., 2020). Thus, to investigate the development and treatment of SUD using preclinical animal models more effectively, choice (type, frequency, and dose) and homecage environment should be taken into consideration. Likewise, procedures that maximize the potential for development and expression of individual differences in intake/preference have the ability to provide previously under-appreciated data types/parameters and predictive factors related to substance use and SUD.

One potential solution that supports both voluntary intake and social housing is the use of a home cage monitoring (HCM) system. HCM reduces animal stressors that could lead to altered behaviors during experimentation, such as movement or researcher intervention (Mingrone et al., 2020; Robinson & Riedel, 2014; Voikar & Gaburro, 2020). This also allows users to study circadian rhythm cycles and animals’ natural behaviors with more accuracy (Robinson & Riedel, 2014). There are a variety of commercial and open source HCM systems on the market, each with unique characteristics and advantages/disadvantages (de Chaumont et al., 2019; Godynyuk et al., 2019; Iman et al., 2021; Lee et al., 2020; Matikainen-Ankney et al., 2019; Melo et al., 2022; Peleh et al., 2019; Redfern et al., 2017; Robinson & Riedel, 2014; Singh et al., 2019). Some HCM systems focus on tracking and analyzing social interaction (de Chaumont et al., 2019; C. J. M. I. Klein et al., 2022; Peleh et al., 2019; Redfern et al., 2017; Singh et al., 2019), while others focus on continuous monitoring of voluntary liquid or food intake (Godynyuk et al., 2019; Lee et al., 2020; Melo et al., 2022; Robinson & Riedel, 2014; Woodard et al., 2020). Critically, there is currently no low cost or open-source solution that allows for group housed animals to self-administer multiple substances in multiple doses in a homecage environment.

Here we detail development of the Socially Integrated Polysubstance (SIP) system, an opensource solution that allows for group housed rodents to self-administer multiple substances and/or doses with continuous monitoring and measurement, including tracking social interaction by overhead video collection, in a homecage environment. We test the utility of these cages by examining preference and intake patterns across four flavors of Kool-Aid in male and female mice housed in groups of three. Results demonstrate individual and sex differences in flavor preference and intake patterns and highlight the ability of these cages to examine polysubstance use across an extended time frame in group housed rodents.

## Materials and Methods

### Animals

All experiments utilized female and male (as determined by genital appearance at weaning) C57Bl/6 mice aged 9-11 weeks of age at time of arrival to VA Puget Sound. Mice were housed by sex in cages of three on a 12:12 light:dark cycle (lights on at 06:00), and were given *ad libitum* food and water. All animal experiments were carried out in accordance with Association for Assessment and Accreditation of Laboratory Animal Care guidelines and were approved by the VA Puget Sound Institutional Animal Care and Use Committees. Mice were acclimated to the housing room for a week following arrival and subsequently handled for an additional week prior to experimental manipulation. To increase rigor and reproducibility, the study included at least two cohorts of mice each run at separate times.

### Device design and construction

The Socially Integrated Polysubstance (SIP) system (Figure 1) consists of four main parts: 1) RFID system - IDspyder (Phenosys, Berlin, Germany), 2) drinking system - Volumetric Drinking Monitor system (VDM, Columbus Instruments, Columbus, Ohio), 3) custom-built home cage with 3d-printed drinking stations, and 4) computer with required software for data acquisition and analysis. A detailed parts list is provided in Table 1. Each IDspyder can include up to 32 radio frequency identification (RFID) sensors and multiple systems can be wired together to increase sensor capacity beyond 32. Each VDM can include up to eight channels (sippers), drawing from 1-8 liquid reservoirs (e.g., up to 8 different fluid types), each capable of delivering 20 ul of fluid per sipper activation. A 20 ul drop of fluid is always present at the end of the VDM sipper and a new 20 ul drop is dispensed only when the current drop is removed. The custom-built home cage is constructed from laser cut plastic parts and is designed to be modular with options for 1-4 drinking stations per wall (up to 12 stations total with current design, but additional stations could be added by increasing the cage size). Figure 2a shows a 4-station setup (4 on one side) and 2b shows the 6-station setup (3 on each side). Each RFID sensor can detect a RFID transponder within ~4 cm, thus we space each drinking station at least that far apart, so each RFID sensor is only activated when an animal enters the drinking station that RFID sensor is part of and is not activated when an animal enters a nearby drinking station. Each drinking station is constructed from 3d-printed plastic with the goal of ensuring close proximity of the RFID sensor and VDM sipper (Figure 2c,d). Laser cut and 3d print files are available at https://github.com/grace3999/SIP_cage_methodspaper2022. The drinking stations connect from the outside and the top of each cage is purposely left open to allow for overhead video collection (Figure 2e). A single computer runs acquisition software provided by Phenosys and Columbus to acquire RFID and VDM data, respectively. These programs are run continuously while the animals are housed in the cage. Custom Python scripts are then used to integrate the RFID and VDM data streams via common timestamps and are available at https://github.com/grace3999/SIP_cage_methodspaper2022. A brief overview of build instructions is included in Table 2.

**Figure 1.**
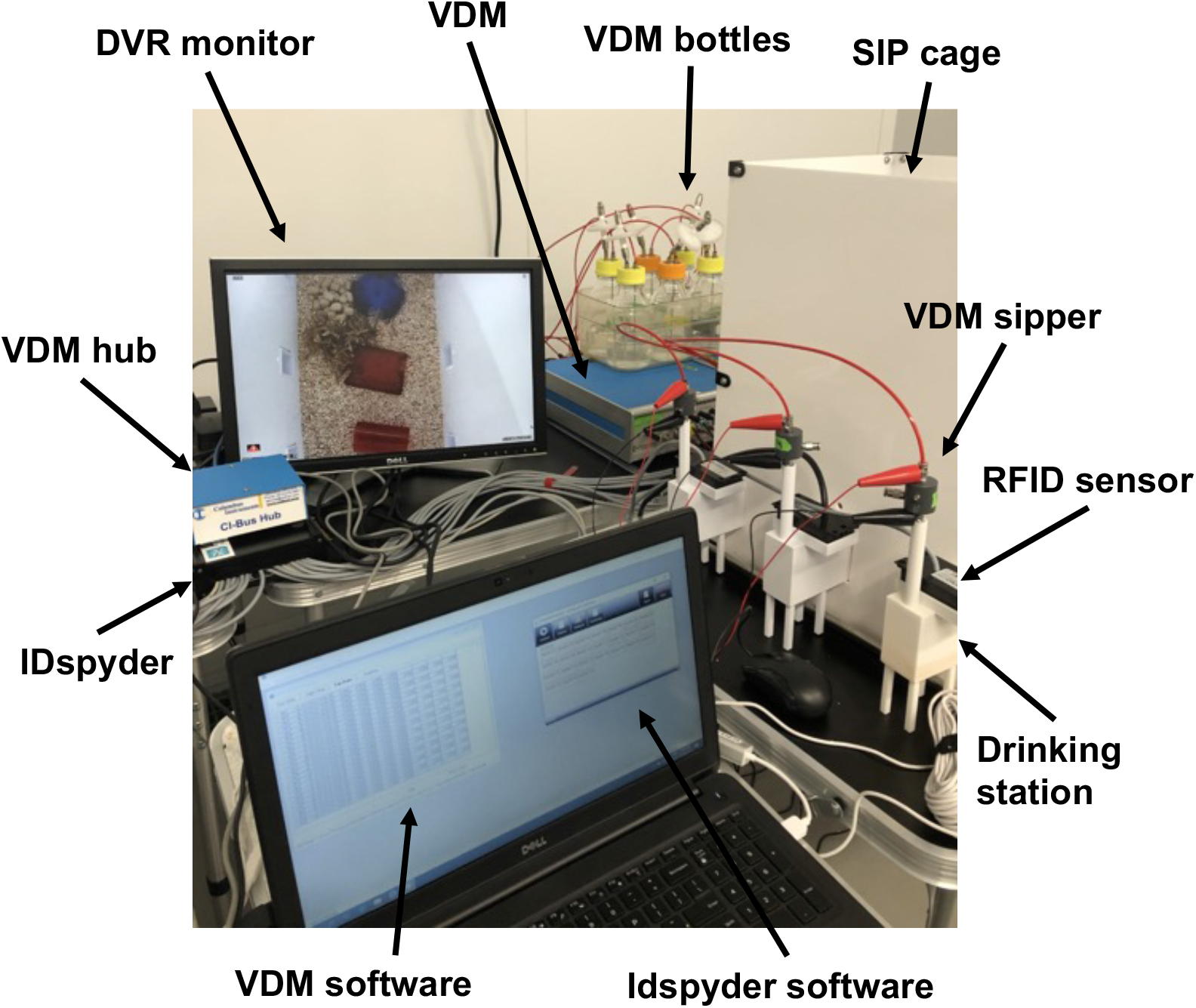
Socially Integrated Polysubstance (SIP) system. Image of assembled SIT system including four main parts: 1) RFID system - IDspyder (base unit and up to 32 sensors, 2) drinking system - Volumetric Drinking Monitor system (VDM; VDM box, bottles, and hub to connect multiple VDM boxes), 3) custom-built homecage with 3d-printed drinking stations, and 4) computer with required software for data acquisition and analysis.

**Figure 2.**
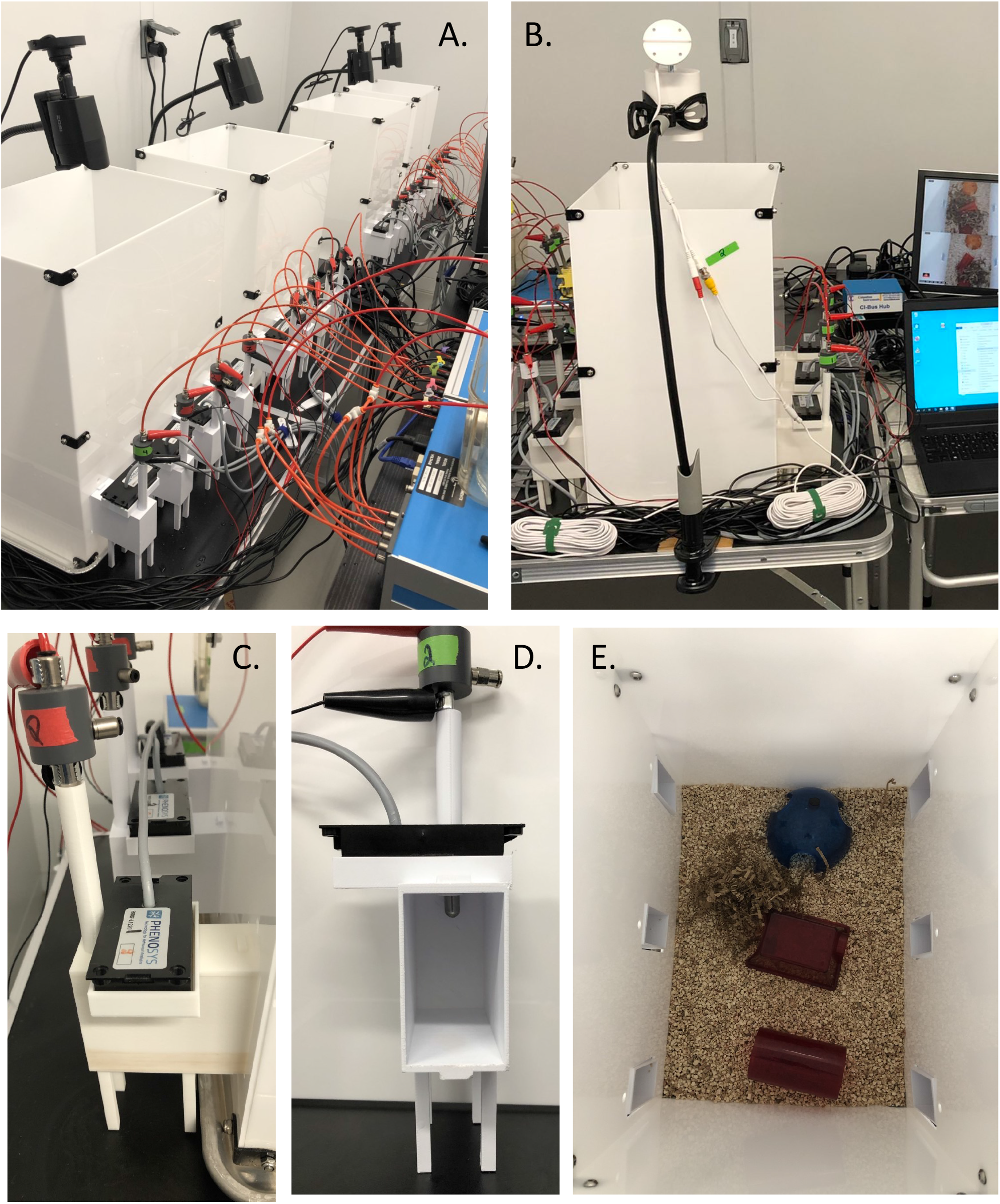
A) SIT 4-station setup (4 on one side), B) SIT 6-station setup (3 on each side). C) Side view of drinking stations connected to SIT wall; each station includes a 3d-printed cubby, VDM sipper, and RFID sensor. D) Front view of drinking station showing access to tip of sipper tube (metal part that extends into the interior) when the animal is inside the drinking station. E) View of SIT 6-station setup from above.

**Table 1:** Parts list

| Component | Quantity | Price | Source |
| --- | --- | --- | --- |
| IDspyder | 1 |  | https://www.phenosys.com/products/rfid_tools/idspyder/ |
| VDM | 1 |  | https://www.colinst.com/products/volumetric-drinking-monitor |
| drinking cubbies | 1-12 |  | 3D printed |
| cage walls | 4 |  | laser cut |
| baker sheet | 1 | 13.88 | https://www.amazon.com/gp/product/B00008UA3Z |
| screws | 24 | 9.49 | https://www.amazon.com/gp/product/B07NVHPPBC |
| hex lock nuts | 24 | 5.23 | https://www.amazon.com/gp/product/B0761189F5 |
| corner brace | 12 | 15.99 | https://www.amazon.com/gp/product/B07FF1CG69 |
| cage bedding |  |  | vivarium |
| cage enrichment |  |  | vivarium |
| mouse chow |  |  | vivarium |

**Table 2:**
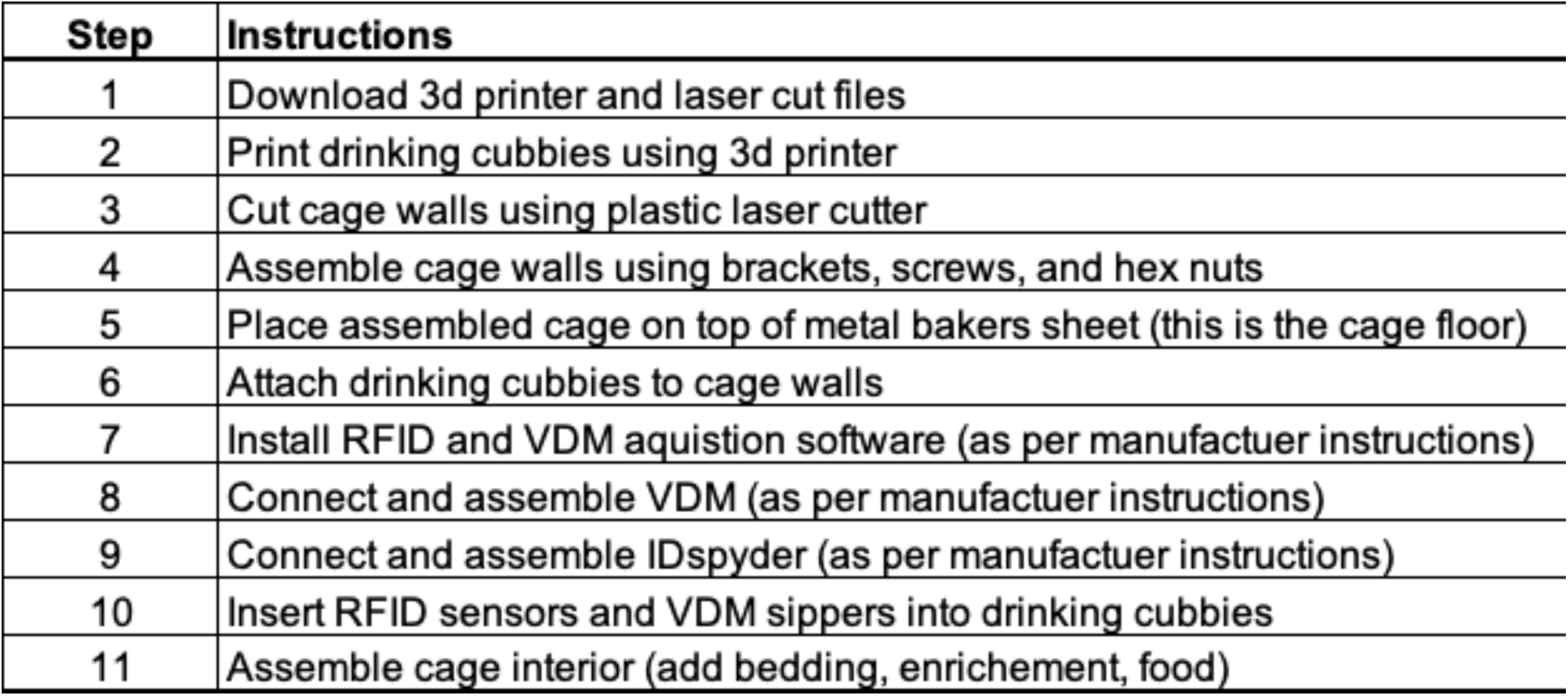
Build instructions

### RFID transponder implantation

Each RFID transponder (Euro I.D., Koln, Germany) is coated in a biocompatible glass material and is 2.12 mm x 12 mm diameter. Prior to implantation, the transponder is sterilized using ethanol and then injected subcutaneously behind the shoulder blades of an anesthetized mice (5% isoflurane) using the provided syringe applicator (Euro I.D., Koln, Germany). On average, the entire process of anesthesia, implantation, and recovery can be completed in three minutes.

### Operating instructions

Following installation of IDspyder and VDM software and build/setup of the SIP cage (Table 2), turn on the IDspyder, VDM, and computer. Start IDspyder acquisition software and load appropriate settings file (this file is customizable and details the number and address of each RFID sensor). Test each RFID sensor using a spare RFID transponder (hold transponder in close proximity to sensor and ensure unique RFID transponder number is displayed in the data acquisition software for the corresponding RFID sensor number/location). Start VDM acquisition software and save new file using a unique and descriptive file name. Test each VDM sipper to ensure fluid is being dispensed and pumps are working appropriately with no leaks (also confirm each sipper is connected to the correct VDM channel). Implant RFID transponder behind shoulder blades of each mouse (under anesthesia). Record weight and unique RFID transponder number for each experimental animal. Place animals in the cage and cover the computer screen with a dark material (computer is left on 24/7 and requires that the screen light be covered to not disrupt the light/dark cycle of the room). Check cages 1x per day and refill food and fluids as needed. When data acquisition is complete, stop programs and save data files (IDspyder saves automatically), prepare metadata files (see examples here: https://github.com/grace3999/SIP_cage_methodspaper2022), and use Python analysis script to combine RFID and VDM datastreams and for subsequent data analysis and visualization.

### Data integration and analysis

Example Jupyter Notebook for data integration can be found at: https://github.com/grace3999/SIP_cage_methodspaper2022. Briefly, this script first cleans the RFID and VDM data (i.e., drops empty rows, fixes datetime, adds light/dark cycle and running time count), adds corresponding metadata (animal demographics and RFID transponder to animal mapping; RFID sensor to fluid type mapping; VDM sipper to fluid type mapping), filters the RFID data by each unique VDM timestamp (e.g., which subject is drinking from what fluid when), and then provides visualization and analysis options. The IDspyder records the duration and starting timestamp of each RFID sensor activation. The VDM records a timestamp for each time a VDM pump is activated (e.g., 20 ul of fluid is removed from the sipper by the animal and another 20 ul is pumped into the sipper). We then find the RFID sensor timestamp that is closest to each unique VDM timestamp and assign the corresponding RFID transponder (i.e., animal) to that unique VDM event. Note, for downstream analysis and visualization, we only include VDM events that occur within 1 second of a RFID event and assume anything outside of 1 second is leakage from the VDM and not an actual animal intake event.

### Kool-Aid flavor preference testing

We tested two separate cohorts of adult female and male C57Bl/6 mice in a set of four SIP cages each equipped with four drinking cubbies (i.e., four liquid options). Four different flavors of Kool-Aid were used, cherry, grape, orange, and blue raspberry, prepared following the provided instructions using sterilized vivarium water. Cohorts were tested for eight days with continuous access to all four flavors in addition to ad libitum food.

### Statistical analysis

Data are expressed as mean ± SEM. Differences between groups were determined using a twotailed Student’s t-test or two-way analysis of variance (ANOVA) followed by posthoc testing using Sidak’s Multiple Comparison. Reported p values denote two-tailed probabilities of p ≤ 0.05 and non-significance (n.s.) indicates p > 0.05. Statistical analysis and visualization were conducted using Graph Pad Prism 4.0 (GraphPad Software, Inc., La Jolla, CA) and with custom Python scripts.

## Results

In our current setup, each SIP cage contains four drinking stations, and each station is equipped with a RFID sensor (connected to IDspyder system) and sipper tube (connected to VDM system). When an animal enters a drinking station, the RFID sensor located at the station is activated and will record the duration of time that sensor is activated (e.g., length of time the animal is in the drinking station). Therefore, using this system, we can measure the duration of each visit and the number of visits to each drinking station.

### RFID system tracks time and duration of visit to individual drinking stations

Figure 3 shows RFID data from 12 male and 12 female adult C57Bl/6 mice recorded from four SIP cages (3 per cage) for 9 days (8 nights) each (2 separate cohorts run with 4 boxes of 3 mice each). Clear light/dark cycle circadian rhythms are apparent across the testing duration (light cycle is shaded in gray). Figure 3a and 3d display data across the whole duration of study; Figure 3b and 3e display the data grouped by hour; Figure 3c and 3f display the data grouped by light/dark cycle. Mice activate (i.e., visit) RFID sensors at an increased number during the dark cycle (t[22]=7.29, p<0.0001), but their time spent at the RFID sensor is longer during the light cycle (t[22]=3.34, p<0.01).

**Figure 3.**
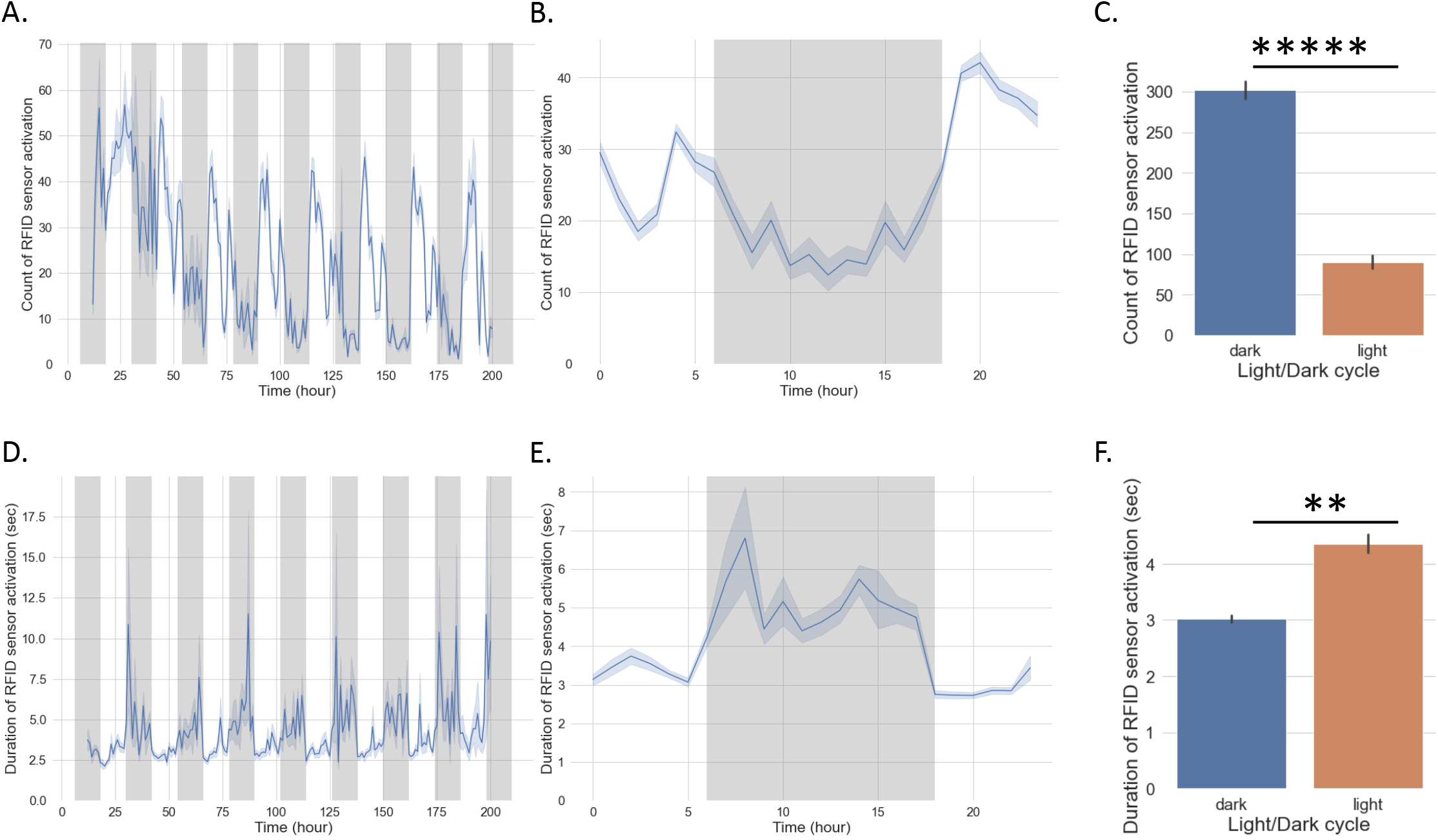
RFID sensor activation data. A-C) RFID sensor count data (number of visits to each sensor). D-F) RFID sensor duration data (average visit duration). A,D: Data per hour across 9 days; B,E: Data averaged per hour; C,F: Data averaged per light/dark cycle. Students t-test. **p ≤ 0.01, ****p ≤ 0.0001. Values represent mean ± SEM.

### VDM system tracks time and volume of drinking events

Figure 4 shows VDM data collected from the same 24 male and female adult C57Bl/6 mice as shown in Figure 2 (4 boxes of 3 mice each run for 9 days with 3 per cage). Clear light/dark cycle circadian rhythms are also present in VDM data across the testing duration (light cycle is shaded in gray). Figure 4a and 4d display data across the whole duration of study; Figure 4b and 4e display the data grouped by hour; Figure 4c and 4f display the data grouped by light/dark cycle. Mice drink from the VDM sippers at an increased number (count) (t[22]=6.96, p<0.0001) and increased amount (volume dispensed per second) (t[22]=8.27, p<0.0001) during the dark cycle.

**Figure 4.**
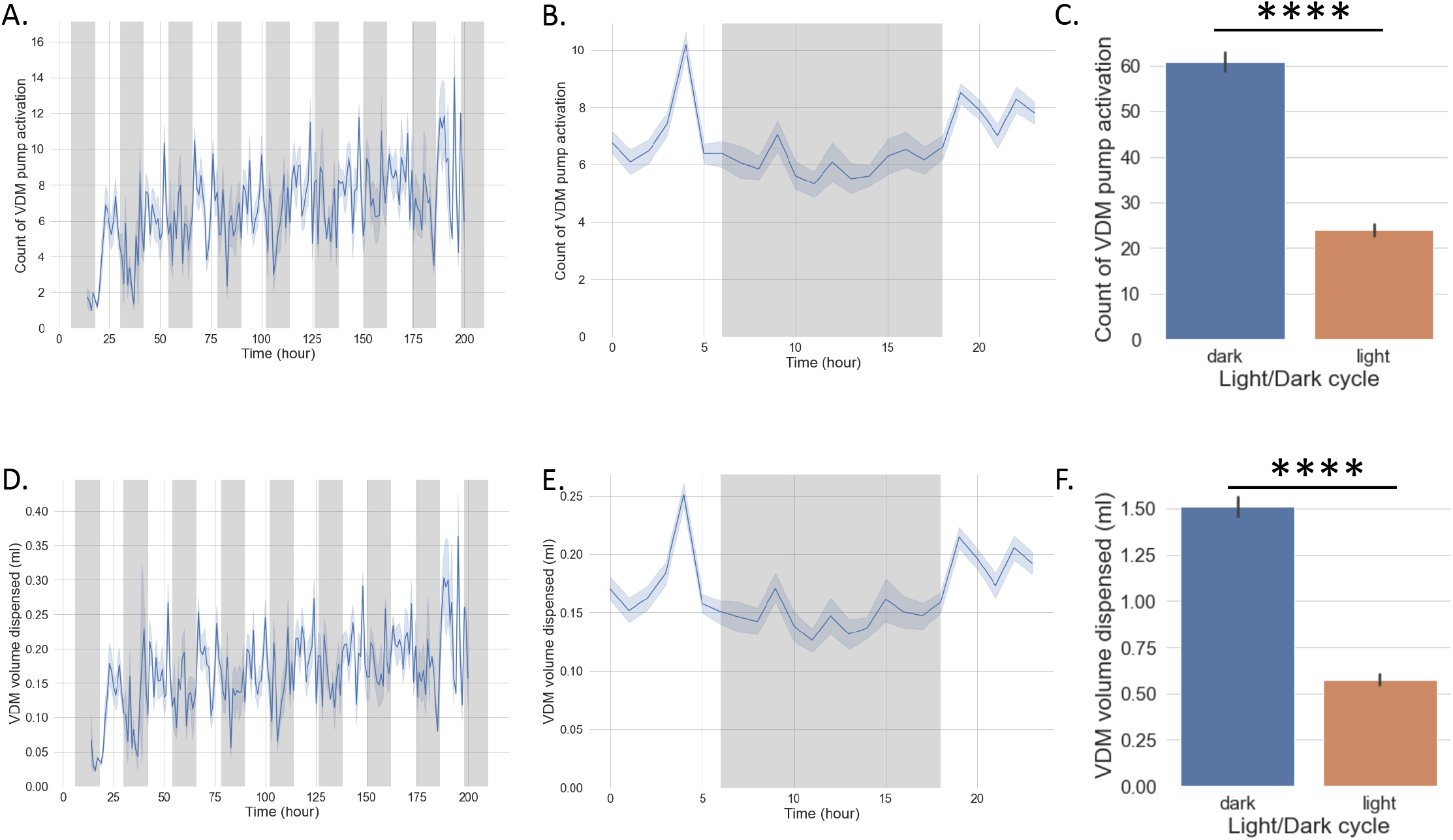
VDM data. A-C) VDM count data (number of times fluid was dispended by VDM). D-F) VDM volume data (volume in ml of fluid dispensed). A,D: Data per hour across 9 days; B,E: Data averaged per hour; C,F: Data averaged per light/dark cycle. Students t-test. ****p ≤ 0.0001. Values represent mean ± SEM.

### Close correspondence between RFID and VDM events

Figure 5 shows how the RFID and VDM timestamps correspond to each other. We computed the difference in time (seconds) between each VDM event and the closest previous RFID event (zero seconds means the VDM event occurred during an RFID event). Figure 5a displays a violin plot of VDM-RFID time difference for each animal, with the majority of events centering at zero. Figure 5b visualizes the VDM-RFID difference distribution for each animal as a kernel density estimate. Likewise, a regression plot of RFID sensor activation duration vs. VDM volume dispensed (Figure 5c) displays a close association between RFID sensor activation and VDM pump activation per animal (r(22) = 0.778, p<0.00001). Finally, Figure 5d shows a raster plot of VDM and RFID timestamps from two example mice across a 24 hour period, note how each VDM mark coincides with an RFID event. Together, these data demonstrate that the vast majority of VDM events occur when the corresponding RFID sensor is also being activated. It is also important to note that not 100% of VDM-RFID differences are zero (likely when the VDM leaks), thus an important data cleaning step is to filter out any VDM events that occur outside of a predefined cutoff threshold (we use 1 second as there can sometime be short a delay between the drinking even and the pump activation).

**Figure 5.**
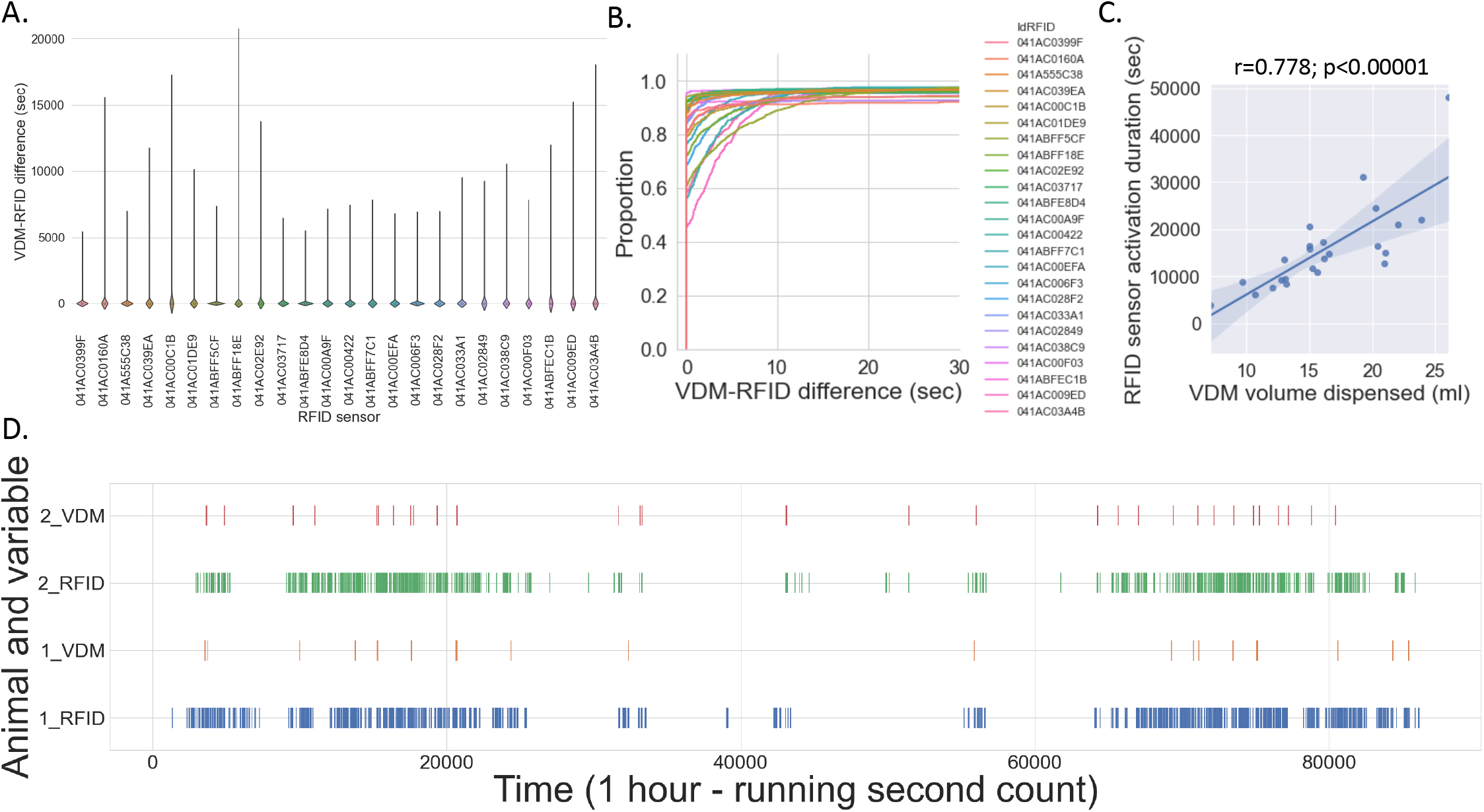
RFID and VDM data are closely aligned. A) Violin plot of VDM-RFID time difference for each animal, note the majority of events are at zero (no difference in VDM and RFID timestamps). B) Distribution of VDM-RFID difference per animal. C) Significant positive correlation between number of RFID sensor activations vs volume of VDM dispended per animal. D) Raster plot of VDM and RFID timestamps from two example mice across a 24 hour period.

### SIP cages reveal sex differences in activity and flavor preference

Figure 6 shows data separated by sex across 24 mice (12 female and 12 male). Female mice visited the drinking stations more frequently than the male mice across all locations (Figure 6a). Likewise, sex differences in flavor preference were also seen, with male mice preferring either the grape or orange, female mice preferring the orange or cherry, and neither sex preferring the blue raspberry (Figure 6b). Individual differences in preference and amount were also seen, with some mice drinking mostly just one flavor and other mice consuming multiple flavors (Figure 6c). Finally Figure 6d shows a raster plot of filtered drinking events for an example mouse, note the different patterns of intake across the four flavors).

**Figure 6.**
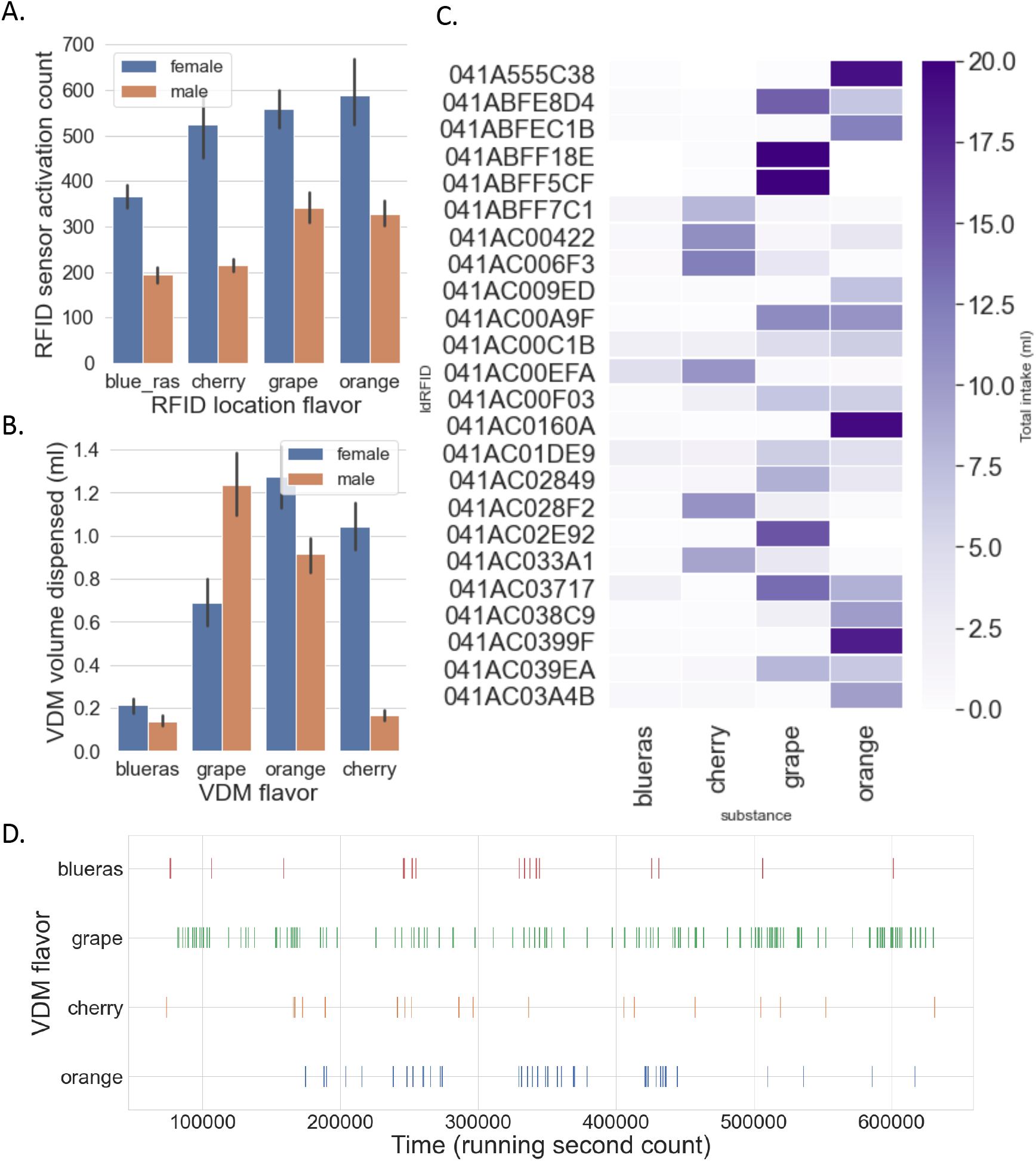
Individual differences in activity and Kool-Aid flavor preference. A) Female mice are more active in the SIT cages than male mice, displaying an increased number of visits to the drinking stations across all locations B) Average intake of 4 Kool-Aid flavors across nine days of access. C) Individual differences in preference and intake. D) Raster plot from representative mouse showing drinking events across multiple days of access to four Kool-Aid flavors.

## Discussion

Choice procedures and the importance of environmental variables are receiving increased attention in the preclinical addiction literature. Polysubstance use, the use of more than one substance either sequentially or simultaneously, has also been recognized as a significant contributor to increased morbidity and mortality in people who use drugs. Critically, existing commercial equipment that allows for social housing and voluntary polysubstance use (e.g., home cage monitoring system) is prohibitively expensive and no open-source solutions exist. With these goals in mind, the current study describes development of the SIP system, which allows 24-hour monitoring of multiple substances in group-housed mice. These cages provide a modular socially interactive environment, have multiple substances measured on a per-second basis without interruption, and permit overhead video collection.

Currently, the IntelliCage (TSE-Systems) is to our knowledge the only available platform (commercial or open source) that allows for both group housing of mice and voluntary access to multiple substances. While the IntelliCage system offers individual recording of long-term and multi-dimensional behavioral patterns for up to 16 mice simultaneously, it can cost upwards of $65,000 USD per cage, limiting the accessibility of this system to only the most well-funded research groups. Furthermore, the IntelliCage does not allow for video monitoring and thus assessment of social interaction is limited. Other commercial options offer either the ability to monitor group-housed animals (e.g., Actual-HCA (Actual Analytics), Digital Ventilated Cage (Techniplast), MultiMouseMonitor (PhenoSys)) or the ability to continuously monitor food or fluid intake (e.g., Phenotyper (Noldus), Chora Feeder (AM Microsystems)). Likewise, open source options are available to either monitor group-housed animals (e.g., LiveMouseTracker (Institute Pasteur)) or monitor food or fluid intake (Home Cage Sipper Device (Creed Lab), DIY-NAMIC (Nautiyal Lab), FED3 (Kravitz Lab)). PiDose (Raymond Lab) offers both a group-housed environment and voluntary intake monitoring of two liquids, but it was designed for daily drug dosing paradigms of a single drug in addition to water access and does not support social interaction monitoring.

Although the SIP system is less expensive than alternatives such as the IntelliCage system, it still uses proprietary hardware and software supplied by the IDspyder and VDM systems. One benefit of using these off-the-shelf solutions for the RFID and intake monitoring parts is the increased ease of assembly and use (no prior knowledge of hardware design, electrical engineering required). To further decrease costs and increase functionality, we are currently working on a version of the SIP system where RFID and intake monitoring are provided by pyControl (Akam et al., 2022), an open source and low cost hardware/software for controlling and monitoring behavioral neuroscience experiments. The upgrade to pyControl will significantly lower costs (no IDsypder or VDM purchase necessary) and add the ability to perform operant based procedures, greatly increasing experimental design capabilities.

Here we present data from the SIP cages when they are equipped with four different flavors of Kool-aid. These data demonstrate the ability of the system to simultaneously monitor voluntary continuous intake of multiple substances in group housed mice, revealing sex and individual differences in Kool-aid flavor preference and activity patterns. Ongoing work is now focused on investigation of single vs. polysubstance paradigms (e.g., multiple doses of alcohol vs. multiple doses of fentanyl vs. multiple doses of alcohol and fentanyl available) in male and female mice and a comparison of intake patterns and preference in single vs. group housed mice. Future plans will incorporate additional substances such as psychostimulants (cocaine, methylphenidate), cannabinoids (THC, CBD), and/or sedative-hypnotics (barbiturate, benzodiazepine). We are also working to develop computer vision algorithms using open source software such as DeepLabCut and plan to integrate RFID and video data (similar to LiveMouseTracker) for downstream analysis of activity/rest patterns and social interaction. Finally, we are developing additional Python scripts for modeling of individual and group differences in intake and preference patterns using unsupervised machine learning and statespace representation.

In conclusion, the SIP cage system is a first step towards designing a rodent model of substance use that more closely resembles the experience of people who use drugs. We envision this system as a tool to help increase understanding of the complex behavioral and pathophysiological mechanisms underlying substance use and development of SUD.

## ABBREVIATIONS

HCM: home cage monitoring
RFID: radio frequency identification
SIP: Socially Integrated Polysubstance
SUD: substance use disorder
VDM: Volumetric Drinking Monitor

## DISCLAIMER

The views expressed in this scientific presentation are those of the author(s) and do not reflect the official policy or position of the U.S. government or Department of Veteran Affairs.

## DECLARATIONS

### Ethics approval and consent to participate

All animal experiments were conducted in accordance with Association for Assessment and Accreditation of Laboratory Animal Care guidelines and were approved by the VA Puget Sound Institutional Animal Care and Use Committee.

### Consent for publication

Not applicable.

### Availability of data and materials

The data in this study are available from the corresponding author upon reasonable request.

### Authors’ contributions

The work presented here was carried out in collaboration among all authors. KL, BB, and AS contributed to conception and design of the study. KL, ZCW, SJL, and AS collected and analyzed data. KW, ZCW, MP, and AS wrote the first draft of the manuscript. All authors contributed to manuscript revision, read, and approved the final manuscript.

## Acknowledgements

We would like to thank Scott Ng Evans, Traci J. Weber, Cindy Pekow, DVM, Kari Koszdin, DVM, and Lena Strait-Bodey for technical assistance and veterinary care. We would also like to thank Val Collins for helpful comments and edits on the manuscript.

